# The histone variant H2A.Z is required to establish normal patterns of H3K27 methylation in *Neurospora crassa*

**DOI:** 10.1101/2020.04.07.029595

**Authors:** Abigail J. Courtney, Masayuki Kamei, Aileen R. Ferraro, Kexin Gai, Qun He, Shinji Honda, Zachary A. Lewis

**Author notes:** Corresponding Author: Zachary A. Lewis, University of Georgia, Athens, GA.

## Abstract

*Neurospora crassa* contains a minimal Polycomb repression system, which provides rich opportunities to explore Polycomb-mediated repression across eukaryotes and enables genetic studies that can be difficult in plant and animal systems. Polycomb Repressive Complex 2 is a multi-subunit complex that deposits mono-, di-, and tri-methyl groups on lysine 27 of histone H3, and tri-methyl H3K27 is a molecular marker of transcriptionally repressed facultative heterochromatin. In mouse embryonic stem cells and multiple plant species, H2A.Z has been found to be co-localized with H3K27 methylation. H2A.Z is required for normal H3K27 methylation in these experimental systems, though the regulatory mechanisms are not well understood. We report here that *Neurospora crassa* mutants lacking H2A.Z or SWR-1, the ATP-dependent histone variant exchanger, exhibit a striking reduction in levels of H3K27 methylation. RNA-sequencing revealed downregulation of *eed*, encoding a subunit of PRC2, in an *hH2Az* mutant compared to wild type and overexpression of EED in a Δ*hH2Az*;Δ*eed* background restored most H3K27 methylation. Reduced *eed* expression leads to region-specific losses of H3K27 methylation suggesting that EED-dependent mechanisms are critical for normal H3K27 methylation at certain regions in the genome.

**AUTHOR SUMMARY:** Eukaryotic DNA is packaged with histone proteins to form a DNA-protein complex called chromatin. Inside the nucleus, chromatin can be assembled into a variety of higher-order structures that profoundly impact gene expression. Polycomb Group proteins are important chromatin regulators that control assembly of a highly condensed form of chromatin. The functions of Polycomb Group proteins are critical for maintaining stable gene repression during development of multicellular organisms, and defects in Polycomb proteins are linked to disease. There is significant interest in elucidating the molecular mechanisms that regulate the activities of Polycomb Group proteins and the assembly of transcriptionally repressed chromatin domains. In this study, we used a model fungus to investigate the regulatory relationship between a histone variant, H2A.Z, and a conserved histone modifying enzyme complex, Polycomb Repressive Complex 2 (PRC2). We found that H2A.Z is required for normal expression of a PRC2 component. Mutants that lack H2A.Z have defects in chromatin structure at some parts of the genome, but not others. Identification of PRC2-target domains that are differentially dependent on EED provides insights into the diverse mechanisms that regulate assembly and maintenance of facultative heterochromatin in a simple model system.

**Data Reference Numbers:** GSE146611

## INTRODUCTION

In eukaryotes, DNA-dependent processes in the nucleus are regulated by chromatin-based mechanisms (1). One heavily studied group of proteins that are particularly important for maintaining stable gene repression are the Polycomb Group (PcG) proteins. In plants and animal cells, PcG proteins assemble into Polycomb Repressive Complexes 1 and 2 (PRC1 and PRC2), which play key roles in repression of developmental genes, as reviewed in (2-6). PRC2 is a multi-subunit complex that deposits mono-, di-, and tri-methyl groups on lysine 27 of histone H3, and tri-methyl H3K27 is a molecular marker of transcriptionally repressed facultative heterochromatin (7-10). PcG proteins are absent from the model yeasts, *Saccharomyces cerevisiae* and *Schizosaccharomyces pombe*, but core PRC2 components have been identified and characterized in several fungi, including *Neurospora crassa, Fusarium graminearum, Cryptococcus neoformans, Epichloë festucae*, and *Fusarium fujikoroi* (11-17). In these fungi, PRC2 is required for repression of key fungal genes suggesting that this enzyme complex is functionally conserved between fungi, plants, and animals (13, 14, 18).

In *N. crassa*, the catalytic subunit of PRC2 is SET-7, a protein with homology to EZH1/EZH2 in humans and curly leaf (CLF), medea (MEA), or swinger (SWN) in *Arabidopsis* (9, 10, 19-25). *N. crassa* EED and SUZ12 are respectively homologous to Drosophila Esc and su(z)12, human EED and SUZ12, and *Arabidopsis* fertilization independent endosperm (FIE; a homolog of EED), and SUZ12 homologs embryonic flower 2 (EMF2), vernalization 2 (VER2), or fertilization independent seed 2 (FIS2) (24, 26, 27). *N. crassa* CAC-3 (also called NPF) is an accessory subunit homologous to mammalian retinoblastoma binding protein 46/48 (RBAP46/68) in humans, and multicopy suppressor of IRA1-5 (MSI1-5) in *Arabidopsis* (28-30). In contrast to PRC2, PRC1 components appear to be absent from the fungal kingdom (31).

The presence of a minimal Polycomb repressive system in well studied fungi such as *N. crassa* provides an opportunity to explore the diversity of Polycomb-mediated repression across eukaryotes and enables genetic studies that can be difficult in plant and animal systems. Indeed, genetic studies have provided insights into PRC2 control in *Neurospora*. Deletion of CAC-3 causes region-specific losses of H3K27me3 at telomere-proximal domains, and telomere repeat sequences are sufficient to nucleate a new domain of H3K27me3-enriched chromatin (14, 32). In constitutive heterochromatin domains, heterochromatin protein-1 (HP1) prevents accumulaton of H3K27me3 (33, 34). Thus, regulation of H3K27 methylation occurs at multiple levels. Despite recent advances, the mechanisms that regulate PRC2 in fungal systems and eukaryotes in general is poorly understood.

In addition to the core histones (H2A, H2B, H3, and H4), eukaryotes also encode non-allelic histone variants. One of the most conserved and extensively studied histone variants is H2A.Z, which is enriched proximal to transcription start sites (TSS) and in vertebrate enhancers (35-42). Functional studies of H2A.Z have linked presence of this variant in nucleosomes to gene activation, gene repression, maintaining chromatin accessibility, and a multitude of other functions (37, 43-50). Notably, H2A.Z has been implicated in the direct regulation of H3K27 methylation in mouse Embryonic Stem Cells (mESCs) and in plants (51-53). In mESCs, there is a strong correlation between the activity of PRC2, enrichment of H3K27me3, and the presence of H2A.Z (54). Colocalization of SUZ12, a subunit of PRC2, and H2A.Z has been found in mESCs at developmentally important genes, such as HOX clusters (39). In addition, H2A.Z is differentially modified its N- and C-terminal tails at bivalent domains that are “poised” for activation or repression upon differentiation (53, 55). N-terminal acetylation (acH2A.Z) or C-terminal ubiquitylation (H2A.Zub) repress or stimulate the action of PRC2 through interactions with the transcriptional activator BRD2 or the PcG protein complex PRC1 (53). It is important to note that functional studies of H2A.Z are challenging because this histone variant is essential for viability in most organisms, including *Drosophila, Tetrahymena*, mouse, and *Xenopus* (56-61).

In *Arabidopsis thaliana*, a genetic interaction between PICKLE (PKL), a chromatin remodeler which promotes H3K27me3, and PIE-1 (homolog to SWR-1), the remodeler which deposits H2A.Z, was recently reported (52). PKL has been found by ChIP-seq at loci enriched for H3K27me3 and is proposed to determine levels of H3K27me3 at repressed genes in *Arabidopsis* (62). In rice callus and seedlings, H2A.Z is found at the 5’ and 3’ ends of genes that are highly expressed. In repressed genes, H2A.Z is found along the gene body, and this pattern closely mimics the presence of H3K27me3 (51). This is a notable difference between plants and other eukaryotes.

We investigated the relationship between H2A.Z and PRC2 in the filamentous ascomycete *Neurospora crassa* and report that H2A.Z is required for normal enrichment of H3K27me2/3 across the genome. Our findings show that loss of H2A.Z leads to region-specific depletion of H3K27me2/3 in *N. crassa*. Expression levels of *eed*, encoding a PRC2 subunit, are reduced in the absence of H2A.Z and ectopic expression of *eed* can restore H3K27me2/3 in an H2A.Z-deficient strain. Together, these data suggest that H2A.Z regulates facultative heterochromatin through transcriptional regulation of the PRC2 component EED and points to differential requirements for EED at discrete PRC2-target domains.

## RESULTS

### Normal patterns of H3K27me2/3 enrichment require the presence of H2A.Z or SWR-1

Normal H3K27me2/3 patterns in plants and in mESCs depend on the histone variant H2A.Z (39, 52), but the underlying mechanism is poorly understood. To determine if H2A.Z also plays a role in Polycomb Group repression in *N. crassa*, we performed ChIP-seq to examine H3K27me2/3 enrichment in an H2A.Z deletion strain (Δ*hH2Az::hph, hereafter ΔhH2Az)* and compared this to wild type. Inspection of the data in the IGV genome browser (63) revealed that the Δ*hH2Az* mutant displayed a significant reduction in H3K27me2/3 (Figure 1A). To quantify the change in H3K27me2/3 patterns, we called peaks of H3K27me2/3 enrichment using Hypergeometric Optimization of Motif EnRichment (HOMER; version 4.8) (64). We identified 325 peaks of H3K27me2/3 in wild type, hereafter referred to as PRC2-target domains (Table S2). Consistent with previous studies, these peaks comprised ∼6% of the *N. crassa* genome (14, 33). These regions are typically larger than single genes, ranging in size from 500 bp to 108 kb, with an average size of 7.7 kb. We next plotted H3K27me2/3 levels across the 5’ end of all 325 domains for wild type and Δ*hH2Az* (Figure 1B). Inspection of heatmaps and the genome browser revealed that H3K27me2/3 levels were reduced in many, but not all PRC2-target domains in Δ*hH2Az*. Using HOMER software to identify PRC2-target domains in Δ*hH2Az* revealed 239 peaks (Table S3). These were slightly smaller, with an average size of 5.5 kb, and comprised only 3% of the *N. crassa* genome. To determine if the peaks observed in the Δ*hH2Az* strain are in wild type locations we only compared peaks from assembled contigs. Using bedtools intersect we found that all peaks in Δ*hH2Az* overlap with wild type peaks, indicating that Δ*hH2Az* exhibits significant loss of H3K27me2/3 from normal domains but does not gain H3K27me2/3 in new locations (Table S4).

**Figure 1:**
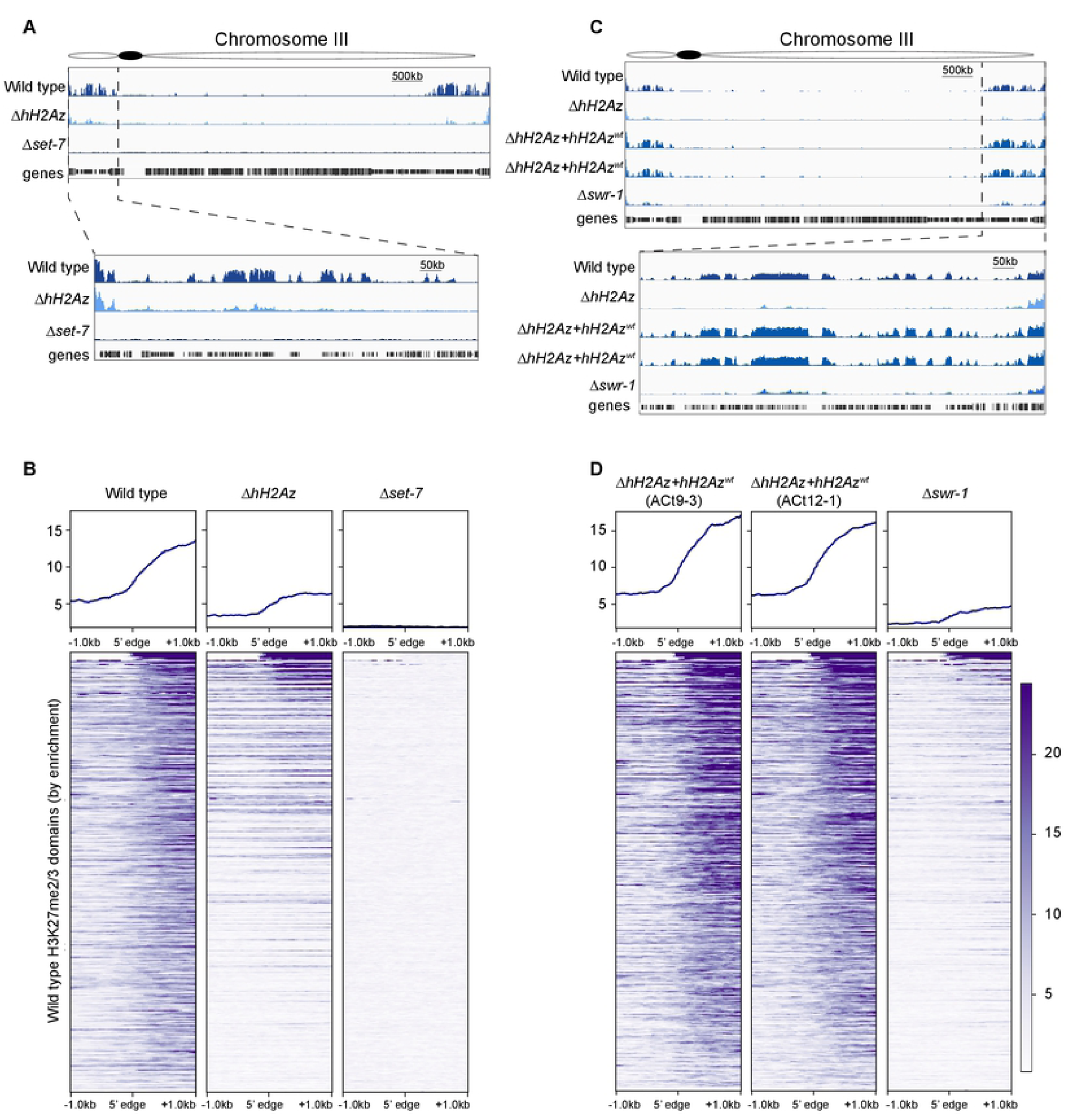
H2A.Z is required for normal patterns of H3K27 methylation. A) Genome browser images illustrate H3K27me2/3 enrichment on *N. crassa* Linkage Group (“chromosome”) III for wild type, Δ*hH2Az*, and Δ*set-7*. A segment of chromosome III is displayed at higher resolution to illustrate depletion of internal H3K27me2/3 domains. B) H3K27me2/3 in the Δ*hH2Az* strain exhibits striking depletion of most H3K27me2/3 domains, with overall lower enrichment of this modification. Heatmaps display 325 PRC2-target domains (rows) ordered by wild type enrichment for wild type, Δ*hH2Az*, and Δ*set-7* strains centered on the 5’ end of each domain + or – 1,000 bp for a total window size of 2000 bp. C) Genome browser images illustrate H3K27me2/3 enrichment on chromosome III for wild type, Δ*hH2Az*, and two ectopic complemented strains of Δ*hH2Az+hH2Az*^wt^, as well as the Δ*swr-1* strain. The segment of chromosome III is displayed at higher resolution to illustrate rescue by complementation and depletion of H3K27me2/3 in Δ*swr-1* background. D) Heatmaps of H3K27me2/3 rescue in complemented strains (Δ*hH2Az+hH2Az*^wt^ [ACt9-3 and ACt12-1]) and depletion in the Δ*swr-1* strain. The heatmaps are ordered as in *B* and depict the domain boundary + or – 1,000 bp for a total window size of 2,000bp.

Since H2A.Z is required for maintaining genome stability in yeast and animals, our findings raised the possibility that a second site mutation could be responsible for the observed phenotype (46, 47, 65, 66). To confirm that loss of H3K27me2/3 was due to the absence of H2A.Z, we first backcrossed the original deletion strain (FGSC 12088) to wild type (67). Four independent Δ*hH2Az* progeny all displayed similar reduction in H3K27me2/3 levels (Figure S1). In addition, the backcrossed Δ*hH2Az* strain displayed slow and variable growth (Figure S2) and was hypersensitive to the DNA damaging agent MMS. This is consistent with previous studies that have demonstrated poor growth of Δ*hH2Az* in *S. cerevisiae* and in *N. crassa* (68, 69).

We next introduced a wild type copy of the *hH2Az* gene with its native promoter into Δ*hH2Az* (Figure S3A). This complemented defects in growth and MMS-sensitivity, and fully restored H3K27 methylation, suggesting loss of H2A.Z was responsible for all observed phenotypes in the deletion mutant (Figure 1C and 1D, S2). Because a specific chromatin remodeling complex, SWR1, exchanges H2A.Z for H2A in plants, yeast and animals (69-73), we next examined H3K27me2/3 in a deletion strain lacking the *N. crassa* homolog of the SWR1 ATPase (Δ*swr-1)*. The *swr-1* mutant displayed a similar reduction in H3K27me2/3 (Figure 1C and D). Together, these data demonstrate that H2A.Z is required for normal H3K27me2/3 in *N. crassa*.

### Deletion of *hH2Az* results in region-specific loss of H3K27me2/3

Visual inspection of the ChIP-seq data revealed losses of H3K27me2/3 from PRC2-target domains located at internal (i.e., non-subtelomeric regions >200kb from the telomere repeats) chromosome sites, but not at telomere-proximal sites (i.e., <200kb from the telomere repeats) (Figure 2A). To quantify this, we inspected ChIP-seq results for H3K27me2/3 for both classes and found retention of H3K27 methylation in telomere-proximal regions with progressive loss in domains farther from chromosome ends. Previously published work showed that a *cac-3* deficient strain has H3K27me2/3 loss which was primarily observed in the telomere-proximal regions (14); *cac-3* encodes an accessory subunit of PRC2 in *N. crassa* homologous to the conserved PRC2 components Msl1-5, NURF55, Rpbp46/48, found in plants, *Drosophila*, and humans, respectively. The phenotype reported here for Δ*hH2Az* appears to be the inverse of the Δ*cac-3* phenotype (Figure 2A).

**Figure 2:**
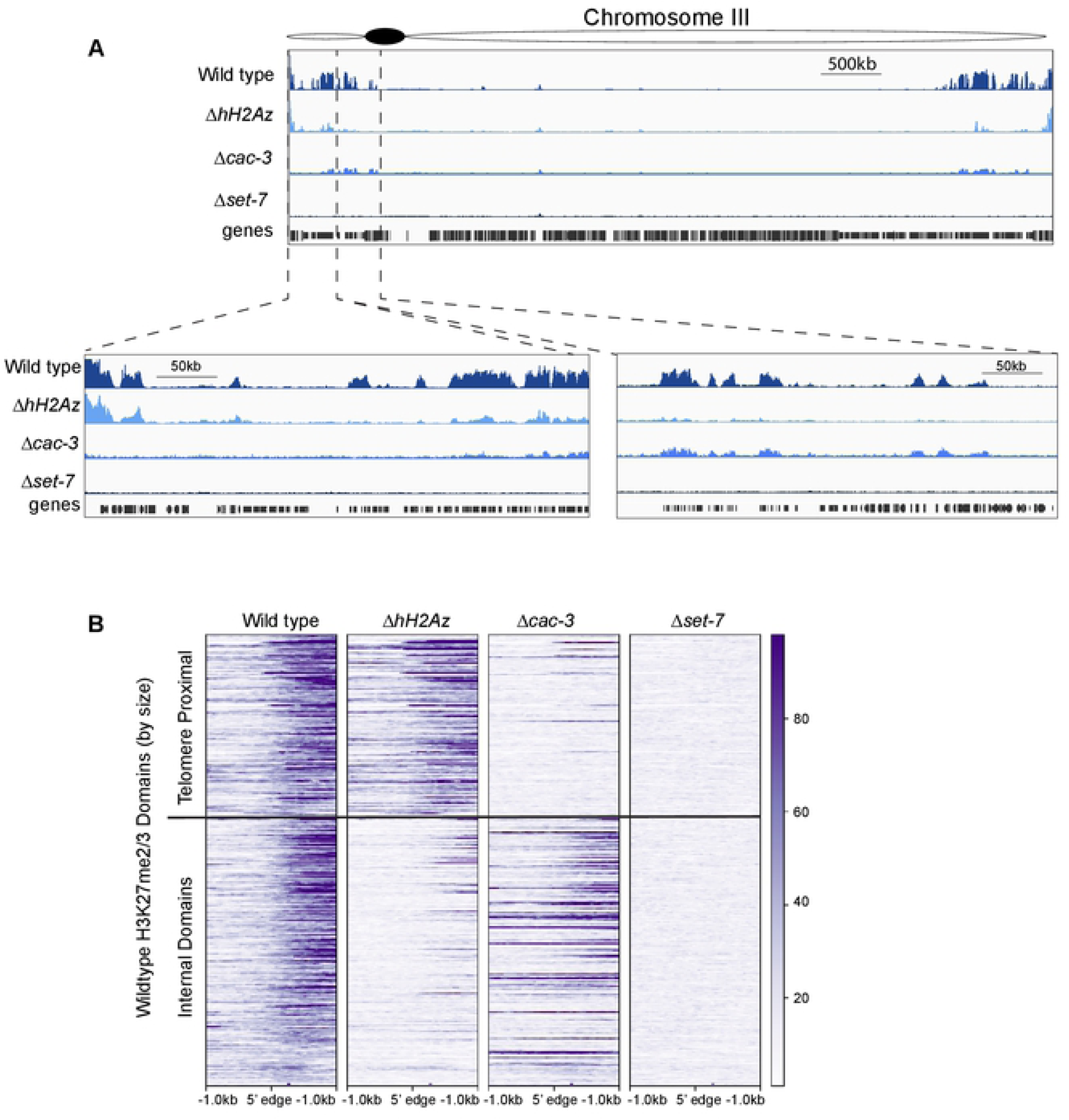
Deletion of *hH2Az* results in region-specific loss of H3K27me2/3. A) Genome browser images illustrate H3K27me2/3 enrichment on Linkage Group (“chromosome”) III for wild type, Δ*hH2Az, Δcac-3*, and Δ*set-7*. The two segments of chromosome III are displayed at higher resolution to visualize region-specific loss. Left panel displays the end of the chromosome to ∼300 kb (left panel) and from ∼400-600 kb (right panel). The telomere-proximal H3K27me2/3 regions are only moderately affected by the deletion of *hH2Az*, whereas internal domains show a more dramatic loss of H3K27me2/3. B) Heatmaps of H3K27me2/3 enrichment for wild type, Δ*hH2Az, Δcac-3*, and Δ*set-7* across PRC2-target domains organized by their proximity to the telomere. The top section is restricted to domains that are <200 kb away from the chromosome ends (“telomere-proximal domains”), plotted from largest to smallest. The bottom of the heatmaps contain the domains that are >200 kb away from chromosome ends (“internal domains”), also plotted from largest to smallest. Heatmaps are centered on the 5’ edge of all 325 PRC2-target domains + or –1,000 bp for a total window size of 2,000 bp. The Δ*hH2Az* strain retains most telomere-proximal H3K27me2/3, as opposed to the Δ*cac-3* strain where almost all H3K27me2/3 enrichment is lost from telomere-proximal regions.

To better visualize which regions of the genome in the Δ*cac-3* or Δ*hH2Az* strains lose enrichment of H3K27me2/3, we again divided all 325 H3K27me2/3 peaks in the wild type strain into telomere-proximal sites (123 peaks, average size 8,261 bp) (Figure 2B, top) and internal sites (186 peaks, average size 7,509 bp) (Figure 2B, bottom). The loss was again most dramatic at the internal regions in the *hH2Az* deletion strain, where most PRC2-target domains showed significant reduction of H3K27me2/3 levels. In contrast, we found that telomere-proximal regions show normal levels of H3K27me2/3.

Previous work demonstrated that the placement of repetitive telomere repeat sequences (5’-TTAGGG-3’) in a euchromatic locus can induce *de novo* H3K27 methylation across large regions (32). Together, these data demonstrate that the absence of H2A.Z is more detrimental for the establishment and/or maintenance of internal domains of H3K27me2/3 in *N. crassa*.

### *Neurospora* H2A.Z localizes to promoter regions but not to PRC2-target domains

We next asked if H2A.Z co-localizes with H3K27 methylation, as has been reported for plants and mESCs (39, 51, 52). We used a strain expressing a C-terminal H2A.Z-GFP fusion protein to perform ChIP-seq with antibodies against H3K27me2/3 and GFP. Visual inspection of the enrichment profiles in a genome browser revealed a mostly mutually exclusive localization pattern (Figure 3A). There are some small H2A.Z peaks that are found in PRC2 target domains, such as in Figure 3A; however, these were rare (Figure 3B). The genomic locations with the highest enrichment for H2A.Z-GFP are the regions immediately before and after the TSS of most genes, with low enrichment in gene bodies and 3’ ends (Figure 3C). On average we find little enrichment of H2A.Z-GFP in the promoters and gene bodies of H3K27me2/3 enriched genes or at the center of H3K27me2/3 peaks, confirming that H3K27me2/3 and H2A.Z are largely mutually exclusive (Figure 3D and 3E).

**Figure 3:**
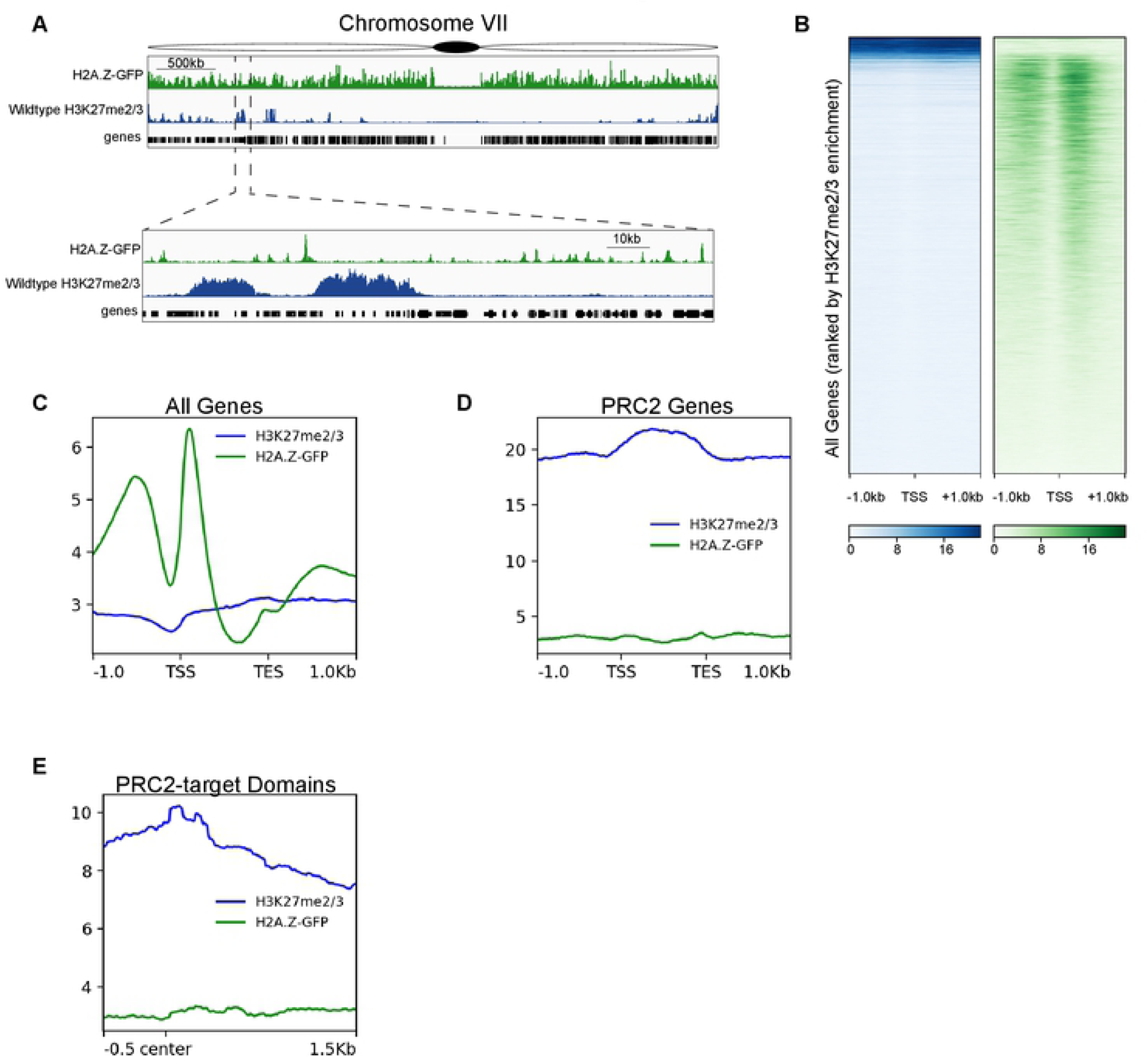
H3K27me2/3 and H2A.Z are not colocalized in *N. crassa* mycelium. A) Genome browser images of ChIP-seq for H2A.Z-GFP (green) and H3K27me2/3 (blue) enrichment across Linkage Group (“chromosome”) VII. A segment of chromosome VII is displayed at higher resolution to visualize the distinct patterns of each modification. Distinct peaks are located at the start of many genes in the genome yet few H2A.Z peaks are present within transcriptionally silent PRC2-target domains. B) Heatmaps of H3K27me2/3 (blue) and H2A.Z-GFP (green) enrichment ordered by H3K27me2/3 enrichment. Heatmaps are centered on the transcription start site (TSS), + or – 1,000 bp for a full window size of 2,000 bp. C) Gene profile of H2A.Z-GFP (green line) and H3K27me2/3 (blue line) enrichment for all genes (fit to 1000 bp for gene body length) in the genome – 1,000 bp upstream of TSS and + 1,000 bp downstream of TES. D) Gene profile of H2A.Z-GFP (green line) and H3K27me2/3 (blue line) enrichment for only H3K27me2/3 enriched genes in the genome – 1,000 bp upstream of TSS and + 1,000 bp downstream of TES. E) Line plot centered on 325 PRC2-target domains displays very low enrichment for H2A.Z-GFP (green line) and high enrichment for H3K27me2/3 (blue line).

To validate H2A.Z enrichment, we also performed ChIP-seq on wild type using an antibody raised against the native *N. crassa* H2A.Z protein (69). These H2A.Z ChIP-seq experiments show the same localization as the H2A.Z-GFP ChIP-seq experiments (Figure S4). The localization of H2A.Z at the TSS of 5,704 genes (over half of all genes) is similar to findings in multiple other organisms (35-42).

### H2A.Z is crucial for proper regulation of one third of the genes in *N. crassa*, including *eed*

Previous studies have implicated H2A.Z in multiple roles related to transcription including gene activation and repression (39, 47, 50, 53, 70, 71). We therefore asked if H2A.Z regulates H3K27me2/3 by regulating expression of one or more PRC2 components. We performed RNA sequencing of wild type, Δ*hH2Az, Δset-7*, and the double mutant Δ*hH2Az;*Δ*set-7* to determine which genes exhibit differential expression in the absence of H2A.Z. Deletion of histone variant H2A.Z causes both positive and negative mis-regulation of a large number of genes (Figure 4A). After Benjamini-Hochberg correction (72), there are 3,308 genes with differential transcription (adjusted *p* value < 0.05). Of these 3,308 genes, there are similar numbers of genes up- and downregulated in the absence of H2A.Z (1,665 genes upregulated and 1,643 downregulated) (Figure 4A, Table S6).

**Figure 4:**
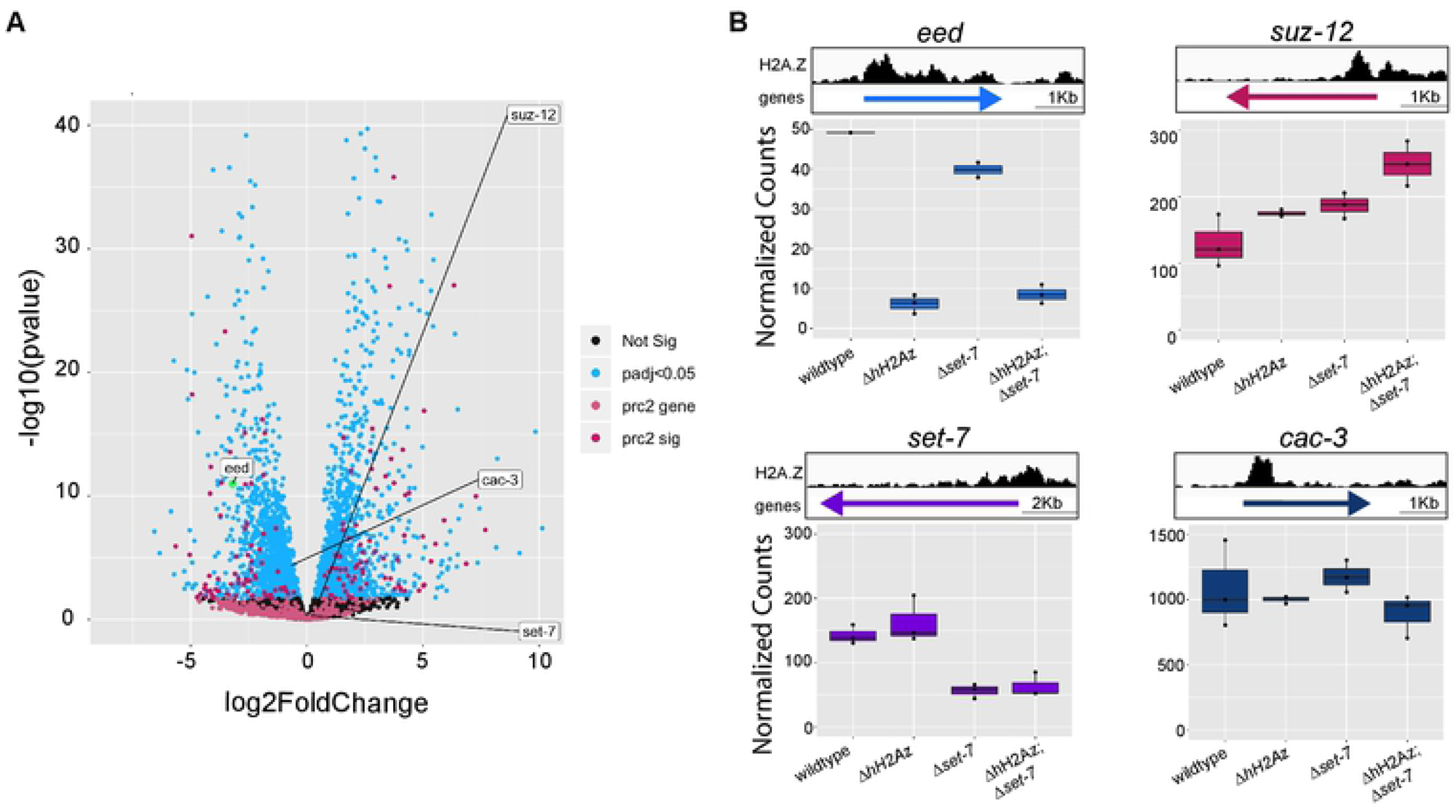
H2A.Z is important for the proper regulation of a large number of genes in *N. crassa*, including *eed*. A) Volcano plot of differentially expressed genes in Δ*hH2Az*. Deletion of *hH2Az* misregulates a large number of genes in both directions; however, there are slightly more genes that are upregulated upon deletion of *hH2Az*. H2A.Z is necessary for the proper expression of a large percentage of genes in *N. crassa*. All members of the PRC2 complex are labeled with text boxes on the plot. Genes enriched for H3K27me2/3 are different shades of pink corresponding to their significance values. The *eed* gene is significantly downregulated in the deletion strain. B) Genome browser images of each gene and its corresponding H2A.Z enrichment, for all PRC2 components, there is enrichment of H2A.Z near the TSS. Boxplots of normalized transcript counts for all subunits of PRC2 (*eed* [light blue], *set-7* [purple], *suz-12* [pink] *cac-3* [dark blue]) in wild type, Δ*hH2Az*, Δ*set-7*, and Δ*hH2Az;*Δ*set-7* backgrounds. Downregulation of *eed* is dependent on *hH2Az* deletion.

We next examined expression levels of genes encoding individual PRC2 components (Figure 4B). We found that expression of *eed* is significantly reduced in Δ*hH2Az* by more than 9-fold (FDR-corrected *p* value = 2.70 x 10^10^), whereas *cac-3, suz-12*, and *set-7* were expressed at similar levels in both wild type and Δ*hH2Az* (Figures 4A and 4B).

The *eed* gene showed the most dramatic change in expression compared to wild type in either Δ*hH2Az* or Δ*hH2Az;*Δ*set-7*, but is expressed normally in the single mutant Δ*set-7*. This indicated that deletion of H2A.Z is likely responsible for its downregulation. As an essential component of PRC2, EED is required for catalytic activity. EED is also important for recognition of the H3K27me2/3 mark and has been implicated in maintenance and/or spreading of H3K27me3 from nucleation sites (73, 74). Since H2A.Z is localized proximal to the promoters of a little over half the genes (5,704) in the *N. crassa* genome, we examined the H2A.Z localization at the *eed* gene. There is a large peak of H2A.Z enrichment at the promoter of *eed* (Figure 4B), which appears to be crucial for normal *eed* expression. Promoters of other PRC2 components are also enriched for H2A.Z, but apparently are not dependent on H2A.Z for their expression. Together, these data suggest that H2A.Z is required for the proper expression of *eed*.

### Overexpression of EED rescues H3K27 methylation levels in the absence of H2A.Z

To determine if downregulation of *eed* is responsible for the depletion of H3K27me2/3 observed in Δ*hH2Az*, we constructed a strain which lacks both *eed* and *hH2Az*, and we introduced a 3x*flag-eed* construct into the *his-3* locus driven by the strong constitutive *clock controlled gene-1/glucose-repressible gene-1* (*ccg-1/grg-1*) promoter *(his-3::Pccg1-3xflag-eed)*.

We calculated expected expression levels of this construct using native *ccg-1* levels, and we expect *eed* to be expressed at approximately 100 times the native level. To confirm this construct was being expressed at the same level in both the Δ*eed* and Δ*eed;ΔhH2Az* backgrounds, we performed an anti-FLAG western blot (Figure S3B). Our results confirm that the deletion of H2A.Z does not alter 3xFLAG-EED expression driven by the *ccg-1* promoter. After performing H3K27me2/3 ChIP-seq in this strain, we find that the majority of H3K27me2/3 peaks are recovered in the genome (Figure 5A), but the growth rate of the Δ*hH2Az* strain is not rescued.

**Figure 5:**
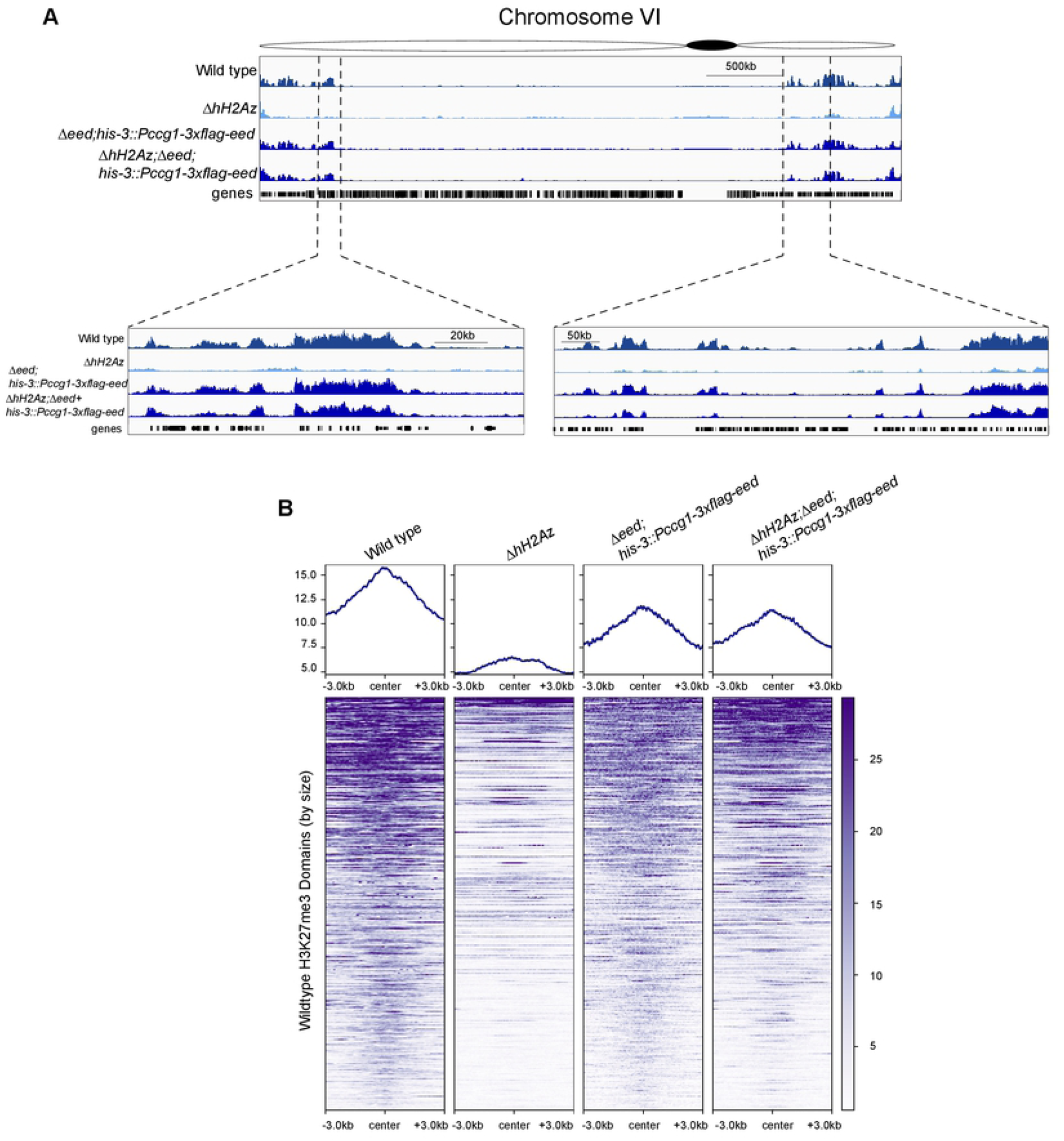
Overexpression of *eed* rescues H3K27 methylation levels in the absence of H2A.Z. A) Partial restoration of H3K27 methylation throughout the genome in a Δ*hH2Az*;Δ*eed* strain containing *his-3::Pccg-1-3xFlag-eed* and overexpressing *eed* at ∼100x the native level. Most H3K27 methylation is restored, though there are some qualitative differences in the peak patterns between the overexpression strain and wild type. B) Heatmaps of H3K27me3 enrichment across 325 PRC2-target domains sorted by size (largest to smallest) centered on each domain + or – 3,000 bp for a full window size of 6,000 bp for wild type, Δ*hH2Az*, Δ*eed;his-3::Pccg-1-3xflag-eed*, and Δ*eed;*Δ*hH2Az;his-3::Pccg-1-3xflag-eed*. Not all domains are fully rescued to wild type levels.

There are some qualitative differences in peak shape and not all peaks are fully restored (Figure 5B), which could indicate that H2A.Z contributes to normal H3K27me2/3 via additional mechanisms. Nevertheless, the significant restoration of H3K27me2/3 suggests that reduced *eed* expression is the major contributor to the loss of H3K27me2/3 in the Δ*hH2Az* strain.

## DISCUSSION

H2A.Z is a highly conserved histone variant that has been linked to gene activation and repression, and control of H3K27 methylation. We report here that *N. crassa* H2A.Z is required for normal methylation of H3K27 in facultative heterochromatin domains. In contrast to the situation in plants and animals, we find that *N. crassa* H2A.Z does not colocalize with H3K27me2/3. In undifferentiated mammalian cells and in plant cells, H2A.Z colocalized with PRC2 components, H3K27me3, SUZ12 or both (39, 51-55). In mESCs, H2A.Z is found at developmentally important loci where SUZ12 is also enriched (39, 55). In addition, this histone variant is proposed to regulate lineage commitment by functioning as a “molecular rheostat” to drive either activation or repression of genes (51, 53, 75). This colocalization of PRC2 and H2A.Z is not seen in differentiated murine cells, and ubiquitylated residues on the C-terminal tail of H2A.Z have been hypothesized as integral for cells to maintain undifferentiated status (53, 55). In plants, H2A.Z displays significant co-localization with H3K27me3 in the gene bodies of PcG-repressed genes even in differentiated tissues (51). Our work highlights an important structural difference between facultative heterochromatin in plants and filamentous fungi. Although we did not observe co-localization of H2A.Z and H3K27me2/3 in *N. crassa*, it remains possible that these two chromatin features overlap in specific developmental cell types (e.g. during sexual development or meiosis). Future work is needed to test this possibility.

In *N. crassa* we generally find histone H2A.Z at the promoters of a large number of genes in the genome. When viewing the localization using a metaplot, which averages the enrichment of all H2A.Z marked nucleosomes, it appears that H2A.Z flanks the TSS. Genome-wide localization of H2A.Z has been performed in a variety of organisms including *Arabidopsis, C. elegans, S. cerevisiae*, mouse, and *Drosophila*. H2A.Z is generally found in the promoters of active and inactive genes, as well as at in vertebrate enhancers (35, 37-42). The +1 nucleosome, first nucleosome after the TSS, containing H2A.Z has been postulated as a lower energy barrier to transcription elongation in *Drosophila* and *Arabidopsis* (35, 36). Our data are consistent with an important promoter-specific role for *N. crassa* H2A.Z.

Indeed, in *N. crassa* we find that the *eed* gene contains a large peak of H2A.Z in the +1 nucleosome, and we find that H2A.Z is required for the proper expression of *eed*. To our knowledge this is the first report of H2A.Z specifically regulating the *eed* gene. Previous studies in mESCs demonstrate that appropriate binding of multiple factors to the *eed* promoter are required for the normal expression of *eed* (76, 77). It is possible that there are *N. crassa* transcription factors that bind to DNA sequences associated with the H2A.Z-containing nucleosome. Nucleosomes that contain H2A.Z protect approximately 120 bp of DNA from MNase digestion as opposed to nucleosomes with canonical H2A that protect 147 bp (78). This may leave more sequence available for transcription factor binding between H2A.Z-containing nucleosomes.

We observed that reduced *eed* expression levels leads to region-specific losses of H3K27me2/3, rather than a more general, or global, reduction. In contrast to our work, reduced *Eed* is reported to cause a global decrease in H3K27me3 in mESCs. In these cells, reduced expression of *Eed* was observed following downregulation of *Oct3/4*, which in turn led to a global reduction of H3K27me3, though these studies did not examine genome-wide patterns of H3K27me3 by ChIP-seq as reported here (76, 77). In *N. crassa*, repetitive sequences (e.g., the canonical telomere repeats) are sufficient to induce an artificial H3K27me3 domain when inserted into a locus normally devoid of H3K27me3 (32). It is interesting that even though we also observe the loss of H3K27 methylation throughout much of the genome, regions proximal to the telomeres retained H3K27me2/3. This might suggest that PRC2 is being recruited to the telomeric region and the downregulation of *eed* causes a defect in the propagation of the H3K27me2/3 modification into topologically associated, nearby regions. Another possibility is that the internal domains have a special requirement for EED in spreading, or for the maintenance of H3K27 methylation following DNA replication. Alternatively, EED may interact directly with transcription factors that control assembly of facultative heterochromatin at certain internal domains, while other PRC2-associated proteins may be more important for targeting PRC2 to telomeres. Future studies are needed to distinguish between these possible working models.

## MATERIALS AND METHODS

### Strains and growth media

Strains used in this study are listed in (Table S1). Strains were grown at 32°C in Vogel’s Minimal Medium (VMM) with 1.5% sucrose or glucose for DNA based protocols, and RNA based protocols, respectively (79). Liquid cultures were shaken at 180 rpm. Crosses were performed on Synthetic Crossing (SC) medium in the dark at room temperature (79). Ascospores were collected 14 days after fertilization. To isolate cross progeny, spores were spread on solid VMM plates containing FGS (1X Vogel’s salts, 2% sorbose, 0.1% glucose, 0.1% fructose, and 1.5% agar) and incubated at 65°C for 1 hour as previously described (79), after which spores were picked using a sterile inoculating needle and transferred to agar slants with appropriate medium (typically VMM). To test for sensitivity to DNA damaging agents, 5 µL of a conidial suspension was spotted on VMM containing FGS (1X Vogel’s salts, 2% sorbose, 0.1% glucose, 0.1% fructose, and 1.5% agar) plates containing concentrations of methyl methanesulfonate (Sigma Aldrich cat. # 129925-5g) between 0.010% and 0.03% (w/v).

To construct the N-terminal FLAG-tagged *eed* allele, we amplified the *eed* region with primers, MK #51: GGCGGAGGCGGCGCGATGCAAATTTGTCGGGACCG and MK #52: TTAATTAATGGCGCGTTACTTCCCCCACCGCTGAA (Table S5), from wild type genomic DNA (FGSC 4200). The amplified fragment was cloned into the *Asc*I site of pBM61::CCGp-N-3xFLAG (80) by InFusion cloning (Takara, cat. # 639648). The new plasmid was then digested with *Dra*I and transformed into a *his-3;mus-52::bar* strain. Primary transformants were selected on VMM plates, and then back-crossed to wild type to isolate homokaryons (*his-3::Pccg-1-3xflag-eed*). We next crossed the homokaryon (*his-3::Pccg-1-3xflag-eed)* to Δ*eed::hph* (FGSC 14852) to obtain Δ*eed;his-3::Pccg-1-3xflag-eed*. 3xFLAG-EED expression and deletion of *eed* deletion were confirmed by western blots probed with anti-FLAG antibody (Sigma Aldrich, cat. # F1804) and genotyped by PCR with primers, LL #155: TCGCCTCGCTCCAGTCAATGACC and LL #466: TGTGGGCGATTTGAGCGTGC, respectively. The Δ*eed;his-3::Pccg-1-3xflag-eed* strain was then crossed to the Δ*hH2Az::hph* (FGSC 12088) strain to obtain Δ*hH2Az;*Δ*eed;his-3::Pccg-1-3xflag-eed*. 3xFLAG-EED expression and deletion of *eed* were confirmed by western blots with anti-FLAG antibody (Sigma Aldrich, cat. # F1804) and genotyping with *eed* deletion primers (see above). Deletion of *hH2Az* was confirmed by PCR with primers AC #24: GAACAAGCCGATTGCTGTCC and AC #23: TGTATAGAACGCTGCCAAGGA.

For the H2AZ-GFP gene replacement construct, a 1-kb segment including the end of the *hH2Az* coding region was amplified by PCR with primers #1577: CGGAAAGGGCAAGTCGTCTG and #1578: CCTCCGCCTCCGCCTCCGCCGCCTCCGCCAGCCTCCTGAGCCTTGGCCT and a 500-bp segment of the 3’ flanking region was amplified with primers #1579: TGCTATACGAAGTTATGGATCCGAGCTCGCTGCACCGAAAAACTCGACG and #1580: GTGACGAGGGGAGATTGCTC. The cassettes containing the GFP segment and the *hph* gene were amplified using M13 forward and reverse primers from *pGFP::hph::loxP* (80). The three fragments were mixed and then assembled by overlapping PCR with primers #1577 and #1580 above. The cassette was transformed into the Δ*mus-52* strain (FGSC 15968) by electroporation.

### Transformation and complementation assays

Transformations were performed as previously described (81). To carry out ectopic complementation of the Δ*hH2Az::hph* strain, two linear gene fragments were electroporated into the mutant strain. Specifically, the *bar* (confers Basta resistance) was amplified with primers LL #148 CCGTCGACAGAAGATGATATTGAAGGAGC and LL #149 AATTAACCCTCACTAAAGGGAACAAAAGC (82) and the wild type *hH2Az* gene fragment including its native promoter (genomic coordinate 1390154-1393398 of GCA_000182925.2 assembly accession) was amplified with primers AC #27 CCCAATCCTAGAATCCCGTCG and AC #21 TAAAAGAGCTGCTGTCGCACG, and fragments were co-integrated into the Δ*hH2Az::hph* strain, followed by selection of transformants on Basta-containing plates (VMM with 2% sorbose, 0.1% glucose, 0.1% fructose, 1.5% agar, and 200 ug/mL Basta). Transformants were transferred to agar slants and then screened by PCR, and Southern blots with the North2South Biotin Random Prime Labeling and Detection Kit (ThermoFisher cat. #17175) and the wild type *hH2Az* gene fragment used as a probe.

### Race tube assay

Race tubes were prepared with 15 mL of VMM plus 1.5% sucrose and 1.5% agar. Strains were grown on VMM plates with 1.5% sucrose and 1.5% agar for 16 hours before using a 6mm cork borer to extract mycelial agar plugs from the edge of growing hyphae. This plug was used for inoculating each tube at one end. Strains were inoculated in triplicate. Measurements were taken at 9, 23, 47 and 60 hours to determine linear growth rates.

### Protein extraction and western blotting

Strains were grown at 32°C shaken in 18×150mm glass test tubes at 180rpm in 5 mL VMM with 1.5% sucrose. After 16 hours, tissue was harvested using filtration, washed once in phosphate buffered saline (PBS), and suspended in 1 mL of ice-cold protein extraction buffer (50mM HEPES pH 7.5, 150mM NaCl, 0.02% NP-40, 1mM EDTA, 1mM phenylmethylsulfonyl fluoride [PMSF; Sigma, P7626], one tablet Roche cOmplete mini EDTA-free Protease Inhibitor Cocktail [Roche, cat. # 11836170001]). Tissue was subjected to sonication by Diagenode Bioruptor UCD-200 to deliver 22.5 30 second pulses at 4°C. After two rounds of centrifugation at 13,200 rpm for 10 minutes, supernatant was mixed with 2x Laemmli buffer and boiled for 5 minutes. Samples were separated by SDS-polyacrylamide gel electrophoresis (SDS-PAGE) and transferred to polyvinylidene difluoride (PVDF) membranes in Tris-Glycine transfer buffer (25mM Tris, 200mM glycine) containing 20% methanol at constant 100V for 1 hour at 4°C. Membranes were blocked with Tris-buffered saline (TBS; 10 mM Tris, pH 7.5, 150mM NaCl) including 3% milk powder for 1 hour and incubated overnight with anti-FLAG antibody (Sigma Aldrich, cat. # F1804) in TBS plus 3% milk. Detection was performed with horseradish peroxidase-conjugated secondary antibodies and SuperSignal West Femto chemiluminescent substrate (ThermoFisher, cat. # 34094).

### Chromatin immunoprecipitation (ChIP)

To carry out ChIP, conidia were inoculated in 5 mL of liquid VMM plus 1.5% sucrose and grown for 18 hours for wild type and other strains with typical growth rates. Slow growing Δ*hH2Az::hph* strains were grown for 24 hours to isolate cultures at a similar developmental stage. ChIP was performed as described previously (83-85). In brief, mycelia were harvested using filtration and were washed once in PBS prior to cross-linking for 10 minutes in PBS containing 1% formaldehyde on a rotating platform at room temperature. After 10 minutes, the reaction was quenched using 125mM glycine and placed back on the rotating platform for five minutes. Mycelia were harvested again using filtration, washed once with PBS, then resuspended in 600 µl of ChIP lysis buffer (50mM HEPES, pH 7.5, 140mM NaCl, 1mM EDTA, 1% Triton X-100, 0.1% sodium deoxycholate, one tablet Roche cOmplete mini EDTA-free Protease Inhibitor Cocktail (Roche, cat. # 11836170001) in 15 mL conical tubes. Chromatin was sheared by sonication after lysing cell walls with the QSONICA Misonix S-4000 ultrasonic processor (amplitude 10, 30 second processing, one second on, one second off), using the Diagenode Bioruptor UCD-200 (Intensity level: Medium, three rounds of 15 minutes (30 seconds on, 30 seconds off) to deliver 22.5 30 second pulses at 4°C. Water temperature was kept at a constant 4°C by using a Biorad cooling module (cat. # 170-3654) with variable speed pump to circulate 4°C water while processing samples. Lysates were centrifuged at 13,000 rpm in an Eppendorf 5415D microcentrifuge for five minutes at 4°C. For ChIP reactions with antibodies against *N. crassa* H2A.Z, 1 µl, 2.5 µl, or 5 µl of antibody was used (antibody supplied by Dr. Qun He, China Agricultural University). For detection of H3K27 di- and tri-methylation (H3K27me2me3; Active Motif 39535), and GFP-tagged H2A.Z (GFP; Rockland 600-301-215) 1 µl of the relevant antibody was used. Protein A/G beads (20 µl) (Santa Cruz, cat. # sc-2003) were added to each sample. Following overnight incubation, beads were washed twice with 1 mL lysis buffer without protease inhibitors, once with lysis buffer containing 500mM NaCl, once with lysis buffer containing 50mM LiCl, and finally with TE (10mM Tris-HCl, 1mM EDTA). Bound chromatin was eluted in TES (50mM Tris pH 8.0, 10mM EDTA, 10% SDS) at 65°C for 10 minutes. Chromatin was de-crosslinked overnight at 65°C. The DNA was treated with RNase A for two hours at 50°C, then with proteinase K for two hours at 50°C and extracted using phenol-chloroform-isoamyl alcohol (25:24:1) followed by chloroform extraction. DNA pellets were washed with 70% ethanol and resuspended in TE buffer. Samples were then prepared for Illumina sequencing.

### RNA extraction

Conidia were inoculated into 100 x 15mm plates containing 25 mL of VMM + 1.5% glucose and grown for 36-48 hours to generate mycelial mats. Using a 9mm cork borer, 5-7 disks were cut out of the mycelial mat and transferred to 125 mL flasks with 50 mL of VMM + 1.5% glucose and allowed to grow for 12 hours at 29°C in constant light while agitating at ∼90-100 rpm. Disks were harvested using filtration and flash frozen with liquid nitrogen. Frozen tissue was transferred to 1.5 mL RNase-free tubes with 100 µl sterile RNase-free glass beads and vortexed to lyse tissue in phenol:chloroform (5:1) pH 4.5. Three sequential acid phenol:chloroform extractions were performed followed by ethanol precipitation using two volumes of ethanol and 1/10 volume of 3M NaOAc pH 5.2, incubated overnight at −20°C. Samples were centrifuged at 13,2000 rpm in 4°C for 30 minutes and pellets were then washed in RNase-free 70% ethanol, and resuspended in RNase-free water. Samples were quantified using the Invitrogen Qubit 2.0 fluorometer (cat. # Q32866) and RNA quality was checked on a denaturing agarose gel. After quality was verified 10 µg of RNA for each sample was subjected to Turbo DNase treatment (Invitrogen, cat. # AM2238) at 37°C for 30 minutes and then another acid phenol:chloroform extraction was performed to inactivate enzyme and purify the RNA. Samples were subjected to another ethanol precipitation as described above, this time with the addition of 1 µL of RNase-free glycogen (5 mg/mL). Samples were centrifuged at 13,200 rpm in 4°C for 30 minutes and the pellets were washed with RNase-free 70% ethanol, then resuspended in RNase-free water. Quality and quantity were again checked with denaturing gel and with the Invitrogen Qubit 2.0 fluorometer. Samples were then prepared for Illumina sequencing.

### ChIP library preparation

Libraries were constructed as described (83-85). In brief, the NEBNext Ultra II End Repair/dA-tailing Module (cat. # E7546S), NEBNext Ultra II Ligation Module (cat. # E7546) were used to clean and A-tail DNA after which Illumina adapters were ligated. The ligation products were amplified to generate dual-indexed libraries using NEBNext Ultra II Q5 Hot Start HiFi PCR Master Mix (cat. # M0543S). Size selection with magnetic beads was performed after the adapter ligation and PCR steps with Sera-Mag SpeedBeads (cat. # 65152105050250) suspended in a solution of (20mM PEG 8000, 1mM NaCl, 10mM Tris-HCl, 1mM EDTA) (86).

### RNA library preparation

Libraries were prepared according to the Illumina TruSeq mRNA stranded Library Kit (cat. # RS-122-2101). In brief, mRNA selection via polyA tails was performed using RNA purification beads and washed with bead washing buffer. Fragmentation and cleanup were performed enzymatically using the Fragment, Prime, Finish Mix and incubated at 94°C for eight minutes. First strand synthesis using the SuperScript II RT enzyme and First Strand Synthesis Act D Mix was incubated as described and second strand synthesis used the Second Strand Marking Mix with resuspension buffer was incubated for one hour to generate cDNA. The final steps in the library preparation are the same as the above ChIP-seq library preparation with exception of two extra bead cleanup steps: one prior to A-tailing and adapter ligation, two after adapter ligation.

Libraries were pooled and sequenced on a NextSeq500 instrument at the Georgia Genomics and Bioinformatics Core to generate single or paired-end reads.

### Data Analysis

For ChIP-seq data, short reads (<20 bp) and adaptor sequences were removed using TrimGalore (version 0.4.4), cutadapt version 1.14 (87), and Python 2.7.8, with fastqc command (version 0.11.3). Trimmed Illumina reads were aligned to the current *N. crassa* NC12 genome assembly available from NCBI (accession # GCA_000182925.2) using the BWA (version 0.7.15) (88), mem algorithm, which randomly assign multi-mapped reads to a single location. Files were sorted and indexed using SAMtools (version 1.9) (89). To plot the relative distribution of mapped reads, read counts were determined for each 50 bp window across the genome using DeepTools to generate bigwigs (version 3.3.1) (90) with the parameters –normalizeUsing CPM (counts per million) and data were displayed using the Integrated Genome Viewer (63). The Hypergeometric Optimization of Motif EnRichment (HOMER) software package (version 4.8) (64) was used to identify H3K27me3 peaks in wild type and Δ*hH2Az* against input using “findPeaks.pl” with the following parameters: -style histone. Bedtools (version 2.27.1) “intersect” (version 2.26.0) was used to determine the number of peaks that intersect with other peak files. Heatmaps, Spearman correlation matrix (Figure S5) and line plots were constructed with DeepTools (version 3.3.1) (90).

For RNA-seq data, short reads (<20 bp) and adaptor sequences were removed using TrimGalore (version 0.4.4), cutadapt version 1.14 (87), and Python 2.7.8, with fastqc command (version 0.11.3). Trimmed Illumina single-end reads were mapped to the current *N. crassa* NC12 genome assembly using the Hierarchical Indexing for Spliced Alignment of Transcripts 2 (HISAT2: version 2.1.0) (91) with parameters –RNA-strandness R then sorted and indexed using SAMtools (version 1.9) (89). FeatureCounts from Subread (version 1.6.2) (92) was used to generate gene level counts for all RNA bam files. Raw counts were imported into R and differential gene expression analysis was conducted using Bioconductor: DeSeq2 (93). Volcano plot and box plots were generated in R using DeSeq2 and ggplot2 (94).

### Data Deposition

Raw sequence data associated with this paper are available through the NCBI GEO database (accession # GSE146611).

## Acknowledgements

This work was supported by grants from the American Cancer Society (RSG-14-184-01-DMC) and the NIH (R01GM132644) to Z.A.L and the National Science Foundation Graduate Research Fellowship Program Grant (DGE-1443117) to A.J.C. We thank the undergraduate students who contributed to this work JongIn Hwang, Vlad Sirbu, Jacqueline Nutter, and Preston Trevor Neal. We are grateful to Robert J. Schmitz and Christina Ethridge for RNA-seq library support and the Georgia Genomic and Bioinformatics Core for sequencing.

## SUPPLEMENTAL FIGURE LEGENDS

**Figure S1: Δ*hH2Az* replicates demonstrating depletion of H3K27me2/3**

Genome browser images of two wild type progeny (top two tracks), and initial four backcrossed sibling *hH2Az* deletion strains on chromosome V. Segment shown at higher resolution to visualize loss of H3K27me2/3.

**Figure S2: Δ*hH2Az* exhibits a slow growth phenotype and is hypersensitive to MMS**

A) MMS Spot test with increasing concentrations of MMS (5 and 10x more *hH2Az* (S532) conidia was used for growth comparable to wild type on sorbose) and decreasing concentrations of conidia.

B) Linear growth rate from race tubes from Δ*hH2Az+hH2Az*^*wt*^ (ACt9-3), Δ*swr*-1, Δ*hH2Az* (S532), and wild type in triplicate.

C) Image of race tubes growing strain in (B) in triplicate.

**Figure S3: Southern Blot confirming ectoptic integration of *hH2Az* gene fragment into *N. crassa***

A) Southern blot of wild type, Δ*hH2Az*, and two ectopic complemented strains (Δ*hH2Az+hH2Az*^*wt*^). Distinct bands in wild type (left arrow) and Δ*hH2Az*. Band corresponding to Δ*hH2Az* and larger band seen in ectopic complemented strains (right arrows). *hH2A*.*z* integration was also confirmed by PCR.

B) FLAG western blot displaying the same expression level of 3xFLAG-EED in both Δ*eed* and Δ*eed;ΔhH2Az* background. 3xFLAG-EED indicated by black arrow (expected size 77.5kD).

**Figure S4: Increasing *N. crassa* H2A.Z antibody concentration improves ChIP-seq resolution**

A) Genome browser image of increasing amount of H2A.Z antibody (1 µL, 2.5 µL, 5 µL).

B) H2A.Z antibody ChIPs in wild type with increasing amounts of H2A.Z antibody. 5 µL is the optimal amount of the H2A.Z antibody to use for the highest resolution of H2A.Z enriched regions.

**Figure S5: Correlation matrix for ChIP-seq replicates**

Spearman correlation matrix for ChIP-seq replicates used in this study

## SUPPLEMENTAL TABLES

**Table S1: Strains used in this study**

**Table S2: H3K27me2/3 domains determined with HOMER for wild type**

**Table S3: H3K27me2/3 domains determined with HOMER for Δ*hH2Az***

**Table S4: H3K27me2/3 domains in common between ΔhH2Az and wild type**

**Table S5: Oligonucleotides used in this study**

**Table S6: Misregulated genes in Δ*hH2Az***

## References

1. Luger K. Structure and dynamic behavior of nucleosomes. Current Opinion in Genetics & Development. 2003;13(2):127–35.

2. Muller J. Transcriptional silencing by the Polycomb protein in Drosophila embryos. EMBO J. 1995;14(6):1209–20.

3. Hennig L, Derkacheva M. Diversity of Polycomb group complexes in plants: same rules, different players? Trends Genet. 2009;25(9):414–23.

4. Simon JA, Kingston RE. Mechanisms of polycomb gene silencing: knowns and unknowns. Nat Rev Mol Cell Biol. 2009;10(10):697–708.

5. Schuettengruber B, Bourbon HM, Di Croce L, Cavalli G. Genome Regulation by Polycomb and Trithorax: 70 Years and Counting. Cell. 2017;171(1):34–57.

6. Kuroda MI, Kang H, De S, Kassis JA. Dynamic Competition of Polycomb and Trithorax in Transcriptional Programming. Annu Rev Biochem. 2020.

7. Muller J, Hart CM, Francis NJ, Vargas ML, Sengupta A, Wild B, et al. Histone methyltransferase activity of a Drosophila Polycomb group repressor complex. Cell. 2002;111(2):197–208.

8. Cao R, Wang L, Wang H, Xia L, Erdjument-Bromage H, Tempst P, et al. Role of histone H3 lysine 27 methylation in Polycomb-group silencing. Science. 2002;298(5595):1039–43.

9. Czermin B, Melfi R, McCabe D, Seitz V, Imhof A, Pirrotta V. Drosophila enhancer of Zeste/ESC complexes have a histone H3 methyltransferase activity that marks chromosomal polycomb sites. Cell. 2002;111(2):185–96.

10. Kuzmichev A, Nishioka K, Erdjument-Bromage H, Tempst P, Reinberg D. Histone methyltransferase activity associated with a human multiprotein complex containing the Enhancer of Zeste protein. Genes Dev. 2002;16(22):2893–905.

11. Veerappan CS, Avramova Z, Moriyama EN. Evolution of SET-domain protein families in the unicellular and multicellular Ascomycota fungi. BMC Evol Biol. 2008;8:190.

12. Aramayo R, Selker EU. Neurospora crassa, a model system for epigenetics research. Cold Spring Harb Perspect Biol. 2013;5(10):a017921.

13. Connolly LR, Smith KM, Freitag M. The Fusarium graminearum histone H3 K27 methyltransferase KMT6 regulates development and expression of secondary metabolite gene clusters. PLoS Genet. 2013;9(10):e1003916.

14. Jamieson K, Rountree MR, Lewis ZA, Stajich JE, Selker EU. Regional control of histone H3 lysine 27 methylation in Neurospora. Proc Natl Acad Sci U S A. 2013;110(15):6027–32.

15. Chujo T, Scott B. Histone H3K9 and H3K27 methylation regulates fungal alkaloid biosynthesis in a fungal endophyte-plant symbiosis. Mol Microbiol. 2014;92(2):413–34.

16. Schotanus K, Soyer JL, Connolly LR, Grandaubert J, Happel P, Smith KM, et al. Histone modifications rather than the novel regional centromeres of Zymoseptoria tritici distinguish core and accessory chromosomes. Epigenet Chromatin. 2015;8.

17. Studt L, Rosler SM, Burkhardt I, Arndt B, Freitag M, Humpf HU, et al. Knock-down of the methyltransferase Kmt6 relieves H3K27me3 and results in induction of cryptic and otherwise silent secondary metabolite gene clusters in Fusarium fujikuroi. Environ Microbiol. 2016;18(11):4037–54.

18. Dumesic PA, Homer CM, Moresco JJ, Pack LR, Shanle EK, Coyle SM, et al. Product Binding Enforces the Genomic Specificity of a Yeast Polycomb Repressive Complex. uCell. 2015;160(1-2):204–18.

19. Jones RS, Gelbart WM. The Drosophila Polycomb-group gene Enhancer of zeste contains a region with sequence similarity to trithorax. Mol Cell Biol. 1993;13(10):6357–66.

20. Chen H, Rossier C, Antonarakis SE. Cloning of a human homolog of the Drosophila enhancer of zeste gene (EZH2) that maps to chromosome 21q22.2. Genomics. 1996;38(1):30–7.

21. Abel KJ, Brody LC, Valdes JM, Erdos MR, McKinley DR, Castilla LH, et al. Characterization of EZH1, a human homolog of Drosophila Enhancer of zeste near BRCA1. Genomics. 1996;37(2):161–71.

22. Goodrich J, Puangsomlee P, Martin M, Long D, Meyerowitz EM, Coupland G. A Polycomb-group gene regulates homeotic gene expression in Arabidopsis. Nature. 1997;386(6620):44–51.

23. Grossniklaus U, Vielle-Calzada JP, Hoeppner MA, Gagliano WB. Maternal control of embryogenesis by MEDEA, a polycomb group gene in Arabidopsis. Science. 1998;280(5362):446–50.

24. Chanvivattana Y, Bishopp A, Schubert D, Stock C, Moon YH, Sung ZR, et al. Interaction of Polycomb-group proteins controlling flowering in Arabidopsis. Development. 2004;131(21):5263–76.

25. Schwartz YB, Pirrotta V. Polycomb silencing mechanisms and the management of genomic programmes. Nat Rev Genet. 2007;8(1):9–22.

26. Schumacher A, Lichtarge O, Schwartz S, Magnuson T. The murine Polycomb-group gene eed and its human orthologue: functional implications of evolutionary conservation. Genomics. 1998;54(1):79–88.

27. Birve A, Sengupta AK, Beuchle D, Larsson J, Kennison JA, Rasmuson-Lestander A, et al. Su(z)12, a novel Drosophila Polycomb group gene that is conserved in vertebrates and plants. Development. 2001;128(17):3371–9.

28. Derkacheva M, Steinbach Y, Wildhaber T, Mozgova I, Mahrez W, Nanni P, et al. Arabidopsis MSI1 connects LHP1 to PRC2 complexes. EMBO J. 2013;32(14):2073–85.

29. Huang S, Lee WH, Lee EY. A cellular protein that competes with SV40 T antigen for binding to the retinoblastoma gene product. Nature. 1991;350(6314):160–2.

30. Qian YW, Wang YC, Hollingsworth RE, Jr., Jones D, Ling N, Lee EY. A retinoblastoma-binding protein related to a negative regulator of Ras in yeast. Nature. 1993;364(6438):648–52.

31. Lewis ZA. Polycomb Group Systems in Fungi: New Models for Understanding Polycomb Repressive Complex 2. Trends Genet. 2017;33(3):220–31.

32. Jamieson K, McNaught KJ, Ormsby T, Leggett NA, Honda S, Selker EU. Telomere repeats induce domains of H3K27 methylation in Neurospora. Elife. 2018;7.

33. Basenko EY, Sasaki T, Ji LX, Prybol CJ, Burckhardt RM, Schmitz RJ, et al. Genome-wide redistribution of H3K27me3 is linked to genotoxic stress and defective growth. P Natl Acad Sci USA. 2015;112(46):E6339–E48.

34. Jamieson K, Wiles ET, McNaught KJ, Sidoli S, Leggett N, Shao YC, et al. Loss of HP1 causes depletion of H3K27me3 from facultative heterochromatin and gain of H3K27me2 at constitutive heterochromatin. Genome Research. 2016;26(1):97–107.

35. Weber CM, Ramachandran S, Henikoff S. Nucleosomes are context-specific, H2A.Z-modulated barriers to RNA polymerase. Mol Cell. 2014;53(5):819–30.

36. Dai X, Bai Y, Zhao L, Dou X, Liu Y, Wang L, et al. H2A.Z Represses Gene Expression by Modulating Promoter Nucleosome Structure and Enhancer Histone Modifications in Arabidopsis. Mol Plant. 2017;10(10):1274–92.

37. Guillemette B, Bataille AR, Gevry N, Adam M, Blanchette M, Robert F, et al. Variant histone H2A.Z is globally localized to the promoters of inactive yeast genes and regulates nucleosome positioning. Plos Biol. 2005;3(12):2100–10.

38. Barski A, Cuddapah S, Cui K, Roh TY, Schones DE, Wang Z, et al. High-resolution profiling of histone methylations in the human genome. Cell. 2007;129(4):823–37.

39. Creyghton MP, Markoulaki S, Levine SS, Hanna J, Lodato MA, Sha K, et al. H2AZ is enriched at polycomb complex target genes in ES cells and is necessary for lineage commitment. Cell. 2008;135(4):649–61.

40. Bargaje R, Alam MP, Patowary A, Sarkar M, Ali T, Gupta S, et al. Proximity of H2A.Z containing nucleosome to the transcription start site influences gene expression levels in the mammalian liver and brain. Nucleic acids research. 2012;40(18):8965–78.

41. Latorre I, Chesney MA, Garrigues JM, Stempor P, Appert A, Francesconi M, et al. The DREAM complex promotes gene body H2A.Z for target repression. Genes Dev. 2015;29(5):495–500.

42. Gomez-Zambrano A, Merini W, Calonje M. The repressive role of Arabidopsis H2A.Z in transcriptional regulation depends on AtBMI1 activity. Nat Commun. 2019;10(1):2828.

43. Bruce K, Myers FA, Mantouvalou E, Lefevre P, Greaves I, Bonifer C, et al. The replacement histone H2A.Z in a hyperacetylated form is a feature of active genes in the chicken. Nucleic acids research. 2005;33(17):5633–9.

44. Neves LT, Douglass S, Spreafico R, Venkataramanan S, Kress TL, Johnson TL. The histone variant H2A.Z promotes efficient cotranscriptional splicing in S. cerevisiae. Genes & Development. 2017;31(7):702–17.

45. Xu Y, Ayrapetov MK, Xu C, Gursoy-Yuzugullu O, Hu Y, Price BD. Histone H2A.Z controls a critical chromatin remodeling step required for DNA double-strand break repair. Mol Cell. 2012;48(5):723–33.

46. Rangasamy D, Greaves I, Tremethick DJ. RNA interference demonstrates a novel role for H2A.Z in chromosome segregation. Nat Struct Mol Biol. 2004;11(7):650–5.

47. Dhillon N, Oki M, Szyjka SJ, Aparicio OM, Kamakaka RT. H2A.Z functions to regulate progression through the cell cycle. Mol Cell Biol. 2006;26(2):489–501.

48. Meneghini MD, Wu M, Madhani HD. Conserved histone variant H2A.Z protects euchromatin from the ectopic spread of silent heterochromatin. Cell. 2003;112(5):725–36.

49. Adam M, Robert F, Larochelle M, Gaudreau L. H2A.Z is required for global chromatin integrity and for recruitment of RNA polymerase II under specific conditions. Molecular and Cellular Biology. 2001;21(18):6270–9.

50. Hu G, Cui K, Northrup D, Liu C, Wang C, Tang Q, et al. H2A.Z facilitates access of active and repressive complexes to chromatin in embryonic stem cell self-renewal and differentiation. Cell Stem Cell. 2013;12(2):180–92.

51. Zhang K, Xu W, Wang C, Yi X, Zhang W, Su Z. Differential deposition of H2A.Z in combination with histone modifications within related genes in Oryza sativa callus and seedling. Plant J. 2017;89(2):264–77.

52. Carter B, Bishop B, Ho KK, Huang R, Jia W, Zhang H, et al. The Chromatin Remodelers PKL and PIE1 Act in an Epigenetic Pathway That Determines H3K27me3 Homeostasis in Arabidopsis. Plant Cell. 2018;30(6):1337–52.

53. Surface LE, Fields PA, Subramanian V, Behmer R, Udeshi N, Peach SE, et al. H2A.Z.1 Monoubiquitylation Antagonizes BRD2 to Maintain Poised Chromatin in ESCs. Cell Rep. 2016;14(5):1142–55.

54. Wang Y, Long H, Yu J, Dong L, Wassef M, Zhuo B, et al. Histone variants H2A.Z and H3.3 coordinately regulate PRC2-dependent H3K27me3 deposition and gene expression regulation in mES cells. BMC Biol. 2018;16(1):107.

55. Ku M, Jaffe JD, Koche RP, Rheinbay E, Endoh M, Koseki H, et al. H2A.Z landscapes and dual modifications in pluripotent and multipotent stem cells underlie complex genome regulatory functions. Genome biology. 2012;13(10):R85.

56. van Daal A, Elgin SC. A histone variant, H2AvD, is essential in Drosophila melanogaster. Mol Biol Cell. 1992;3(6):593–602.

57. Clarkson MJ, Wells JR, Gibson F, Saint R, Tremethick DJ. Regions of variant histone His2AvD required for Drosophila development. Nature. 1999;399(6737):694–7.

58. Liu X, Li B, Gorovsky Ma. Essential and nonessential histone H2A variants in Tetrahymena thermophila. Mol Cell Biol. 1996;16(8):4305–11.

59. Faast R, Thonglairoam V, Schulz TC, Beall J, Wells JR, Taylor H, et al. Histone variant H2A.Z is required for early mammalian development. Curr Biol. 2001;11(15):1183–7.

60. Iouzalen N, Moreau J, Mechali M. H2A.ZI, a new variant histone expressed during Xenopus early development exhibits several distinct features from the core histone H2A. Nucleic acids research. 1996;24(20):3947–52.

61. Ridgway P, Brown KD, Rangasamy D, Svensson U, Tremethick DJ. Unique residues on the H2A.Z containing nucleosome surface are important for Xenopus laevis development. J Biol Chem. 2004;279(42):43815–20.

62. Zhang H, Bishop B, Ringenberg W, Muir WM, Ogas J. The CHD3 remodeler PICKLE associates with genes enriched for trimethylation of histone H3 lysine 27. Plant Physiol. 2012;159(1):418–32.

63. Thorvaldsdottir H, Robinson JT, Mesirov JP. Integrative Genomics Viewer (IGV): high-performance genomics data visualization and exploration. Brief Bioinform. 2013;14(2):178–92.

64. Heinz S, Benner C, Spann N, Bertolino E, Lin YC, Laslo P, et al. Simple combinations of lineage-determining transcription factors prime cis-regulatory elements required for macrophage and B cell identities. Mol Cell. 2010;38(4):576–89.

65. Krogan NJ, Baetz K, Keogh MC, Datta N, Sawa C, Kwok TCY, et al. Regulation of chromosome stability by the histone H2A variant Htz1, the Swr1 chromatin remodeling complex, and the histone acetyltransferase NuA4. Proc Natl Acad Sci USA. 2004;101(37):13513–8.

66. Greaves IK, Rangasamy D, Ridgway P, Tremethick DJ. H2A.Z contributes to the unique 3D structure of the centromere. Proc Natl Acad Sci U S A. 2007;104(2):525–30.

67. Colot HV, Park G, Turner GE, Ringelberg C, Crew CM, Litvinkova L, et al. A high-throughput gene knockout procedure for Neurospora reveals functions for multiple transcription factors. Proc Natl Acad Sci U S A. 2006;103(27):10352–7.

68. Jackson JD, Gorovsky MA. Histone H2A.Z has a conserved function that is distinct from that of the major H2A sequence variants. Nucleic acids research. 2000;28(19):3811–6.

69. Liu X, Dang Y, Matsu-ura T, He Y, He Q, Hong CI, et al. DNA Replication Is Required for Circadian Clock Function by Regulating Rhythmic Nucleosome Composition. Mol Cell. 2017;67(2):203-13.e4.

70. Kim K, Punj V, Choi J, Heo K, Kim JM, Laird PW, et al. Gene dysregulation by histone variant H2A.Z in bladder cancer. Epigenet Chromatin. 2013;6.

71. Valdes-Mora F, Song JZ, Statham AL, Strbenac D, Robinson MD, Nair SS, et al. Acetylation of H2A.Z is a key epigenetic modification associated with gene deregulation and epigenetic remodeling in cancer. Genome Res. 2012;22(2):307–21.

72. Yoav Benjamini, Hochberg Y. Controlling the False Discovery Rate: A Practical and Powerful Approach to Multiple Testing. Journal of the Royal Statistical Society: Series B (Methodological). 1995;57(1):289–300.

73. Xu C, Bian C, Yang W, Galka M, Ouyang H, Chen C, et al. Binding of different histone marks differentially regulates the activity and specificity of polycomb repressive complex 2 (PRC2). Proc Natl Acad Sci U S A. 2010;107(45):19266–71.

74. Hansen KH, Bracken AP, Pasini D, Dietrich N, Gehani SS, Monrad A, et al. A model for transmission of the H3K27me3 epigenetic mark. Nat Cell Biol. 2008;10(11):1291–300.

75. Subramanian V, Fields PA, Boyer LA. H2A.Z: a molecular rheostat for transcriptional control. F1000Prime Rep. 2015;7:01.

76. Ura H, Usuda M, Kinoshita K, Sun C, Mori K, Akagi T, et al. STAT3 and Oct-3/4 control histone modification through induction of Eed in embryonic stem cells. J Biol Chem. 2008;283(15):9713–23.

77. Ura H, Murakami K, Akagi T, Kinoshita K, Yamaguchi S, Masui S, et al. Eed/Sox2 regulatory loop controls ES cell self-renewal through histone methylation and acetylation. Embo Journal. 2011;30(11):2190–204.

78. Tolstorukov MY, Kharchenko PV, Goldman JA, Kingston RE, Park PJ. Comparative analysis of H2A.Z nucleosome organization in the human and yeast genomes. Genome Res. 2009;19(6):967–77.

79. Davis R, de Serres F. Genetic and microbiological research techniques for Neurospora crassa. Methods in Enzymology. 1970;17:79–143.

80. Honda S, Selker EU. Tools for fungal proteomics: multifunctional neurospora vectors for gene replacement, protein expression and protein purification. Genetics. 2009;182(1):11–23.

81. Margolin BS, Freitag, M., and Selker, E.U. Improved plasmids for gene targeting at the his-3 locus of Neurospora crassa by electroporation. Fungal Genetics Newsletter. 1997;44:2.

82. Avalos J, Geever RF, Case ME. Bialaphos resistance as a dominant selectable marker in Neurospora crassa. Curr Genet. 1989;16(5-6):369–72.

83. Ferraro AR, Lewis ZA. ChIP-Seq Analysis in Neurospora crassa. Methods Mol Biol. 2018;1775:241–50.

84. Seymour M, Ji L, Santos AM, Kamei M, Sasaki T, Basenko EY, et al. Histone H1 Limits DNA Methylation in Neurospora crassa. G3 (Bethesda). 2016;6(7):1879–89.

85. Sasaki T, Lynch KL, Mueller CV, Friedman S, Freitag M, Lewis ZA. Heterochromatin controls gammaH2A localization in Neurospora crassa. Eukaryot Cell. 2014;13(8):990–1000.

86. Rohland N, Reich D. Cost-effective, high-throughput DNA sequencing libraries for multiplexed target capture. Genome Research. 2012;22(5):939–46.

87. Martin M. Cutadapt Removes Adapter Sequences From High-Throughput Sequencing Reads. EMBnet. 2011;17(1):2.

88. Li H, Durbin R. Fast and accurate short read alignment with Burrows-Wheeler transform. Bioinformatics. 2009;25(14):1754–60.

89. Li H, Handsaker B, Wysoker A, Fennell T, Ruan J, Homer N, et al. The Sequence Alignment/Map format and SAMtools. Bioinformatics. 2009;25(16):2078–9.

90. Ramirez F, Ryan DP, Gruning B, Bhardwaj V, Kilpert F, Richter AS, et al. deepTools2: a next generation web server for deep-sequencing data analysis. Nucleic acids research. 2016;44(W1):W160–5.

91. Kim D, Paggi JM, Park C, Bennett C, Salzberg SL. Graph-based genome alignment and genotyping with HISAT2 and HISAT-genotype. Nature biotechnology. 2019;37(8):907–15.

92. Liao Y, Smyth GK, Shi W. The Subread aligner: fast, accurate and scalable read mapping by seed-and-vote. Nucleic acids research. 2013;41(10):e108.

93. Love MI, Huber W, Anders S. Moderated estimation of fold change and dispersion for RNA-seq data with DESeq2. Genome biology. 2014;15(12).

94. Wickham H. Ggplot2 : elegant graphics for data analysis. New York: Springer; 2009. viii, 212 p. p.

